# Individualistic attitudes in iterative Prisoner’s Dilemma undermine evolutionary fitness and may drive cooperative human players to extinction

**DOI:** 10.1101/2023.01.02.522489

**Authors:** Erdem Pulcu

**Affiliations:** University of Oxford, Department of Psychiatry, Computational Psychiatry Lab, Oxford, UK

**Keywords:** Prisoner’s Dilemma, Ultimatum Game, evolutionary game theory, computational biology, interpersonal cooperation, social value orientation

## Abstract

Humans perform inarguably the richest plethora of prosocial behaviors in the animal kingdom. Prosocial behaviors such as interpersonal cooperation and altruistic punishment are important for understanding how humans navigate their social environment. The success and failure of strategies human players device have implications for determining their long-term socio-economic/evolutionary fitness. Following the footsteps of Press and Dyson’s (2012), I implemented their evolutionary game theoretic models developed for the Iterated Prisoner’s Dilemma (a behavioral economic probe of interpersonal cooperation) and analysed human choice behavior in the Ultimatum Game (a behavioral economic probe of altruistic punishment) involving 50 human participants versus computerized opponents with prosocial or individualistic social value orientation. Although the results indicate that it is more likely to break cycles of mutual defection in ecosystems in which humans interact with individualistic opponents, analysis of social-economic fitness at the Markov stationary states suggested that this comes at an evolutionary cost. Overall, human players acted in a significantly more cooperative manner than their opponents, but they failed to overcome extortion from individualistic agents, risking “extinction” in 70% of the cases. These findings demonstrate human players might be short-sighted and social interactive decision strategies they device while adjusting to different types of opponents may not be optimal in the long run.

## Introduction

Humans engage in a variety of prosocial behaviors^1-4^. Sometimes people go out of their way to help another person (e.g. interpersonal cooperation^5-7^) or even sacrifice some part of their resources to stand up against unfairness^8,9^, which is commonly known as altruistic punishment. Behavioural-economic games such as the Prisoner’s Dilemma (PD)^10,11^ and the Ultimatum Game (UG)^12,13^ are well-established methods of probing prosocial behaviors (interpersonal cooperation and altruistic punishment, respectively) in experimental/laboratory settings. In 2012, Press and Dyson put forward an important mathematical model of strategy setting in the Iterated PD (IPD, a social-economic game in which two parties interact over a predefined matrix of payoffs repeatedly, allowing players to learn about opponent strategies in time), by which a player can establish a one-sided claim to an unfair share of rewards^14^. Press and Dyson suggested that players using these “zero-determinant” strategies (ZD) can enforce an advantageous linear relationship over the payoffs of their opponents, meaning that if the opponents are rewarded, they would be rewarded even more. This advantageous linear relationship in payoffs is shown to pose a challenge even for evolutionary opponents (i.e. those that can learn and dynamically adjust their behaviours), who are faced with either giving in to extortion, or increasing their Theory of Mind (ToM) sophistication^15^ in order to break free of a relationship that ultimately distributes rewards unfairly. In support of the latter, Press and Dyson (2012) postulated [in their abstract] that:

> *“Only a player with a theory of mind about his opponent can do better, in which case Iterated Prisoner’s Dilemma is an Ultimatum Game*.*”*

In my opinion, this is a very important aspect of Press and Dyson’s work which had been, to the best of my knowledge, overlooked by the scientific community. It raises the possibility of elegantly bridging two key experimental probes of human social-interactive decision-making. This approach can reveal important insights about evolutionary fitness trajectories of strategies humans use in these [cooperation versus competition] scenarios.

Recently, we showed computational model-based evidence to suggest that human players employ second-order or higher^16^ degrees of ToM sophistication in the UG (e.g. taking account of their opponent’s acceptance probability) while choosing between binary Ultimatums^17^ (Fig. 1A). This was captured by the best-fitting computational model to our data, as we showed that human players compute the expected values of Ultimatums by a multiplicative integration of their potential rewards which were modulated by a nonlinear power utility function accounting for participants’ risk attitude^18^, and the opponent’s inferred acceptance probabilities which were modulated by an exponential-logarithmic weighting function^19^ (Fig. 1B). More importantly, this overarching value-based social decision-making model applied irrespective of the opponent’s social value orientation (SVO, which determines a player’s degree of prosociality, e.g. whether the opponents displayed individualistic or prosocial tendencies)^19-21^. A concrete prediction arising from our earlier findings related to the involvement of higher order ToM processes in social-interactive decision-making is that human players should be able to escape extortion. Earlier work suggested that human players can be subdued by extortion, but this approach also impairs extortionate opponent’s own reward trajectory, therefore does not go unpunished^22^. However, as Press and Dyson demonstrated, the ultimate test of any extortion related prediction demands analysis of interpersonal cooperation/competition not within the confines of an experimental timeline (irrespective of whether the experiment is a quantitative simulation or with human players), but at the equilibrium point of these systems. So far, how well human players can escape extortion in iterative games has not been investigated by implementing Press and Dyson’s algebraic approach. Furthermore, no previous work attempted to unify IPD and UG in a way that would allow a game theoretic analysis of [evolutionary social-economic] fitness associated with human interpersonal cooperation at the stationary state (i.e., the equilibrium point of a competitive system at the finite evolutionary horizon). In the subsequent sections, I demonstrate quantitative evidence in support of Press and Dyson’s original claim, followed by some valuable insights into human interpersonal cooperation.

**Figure 1.**
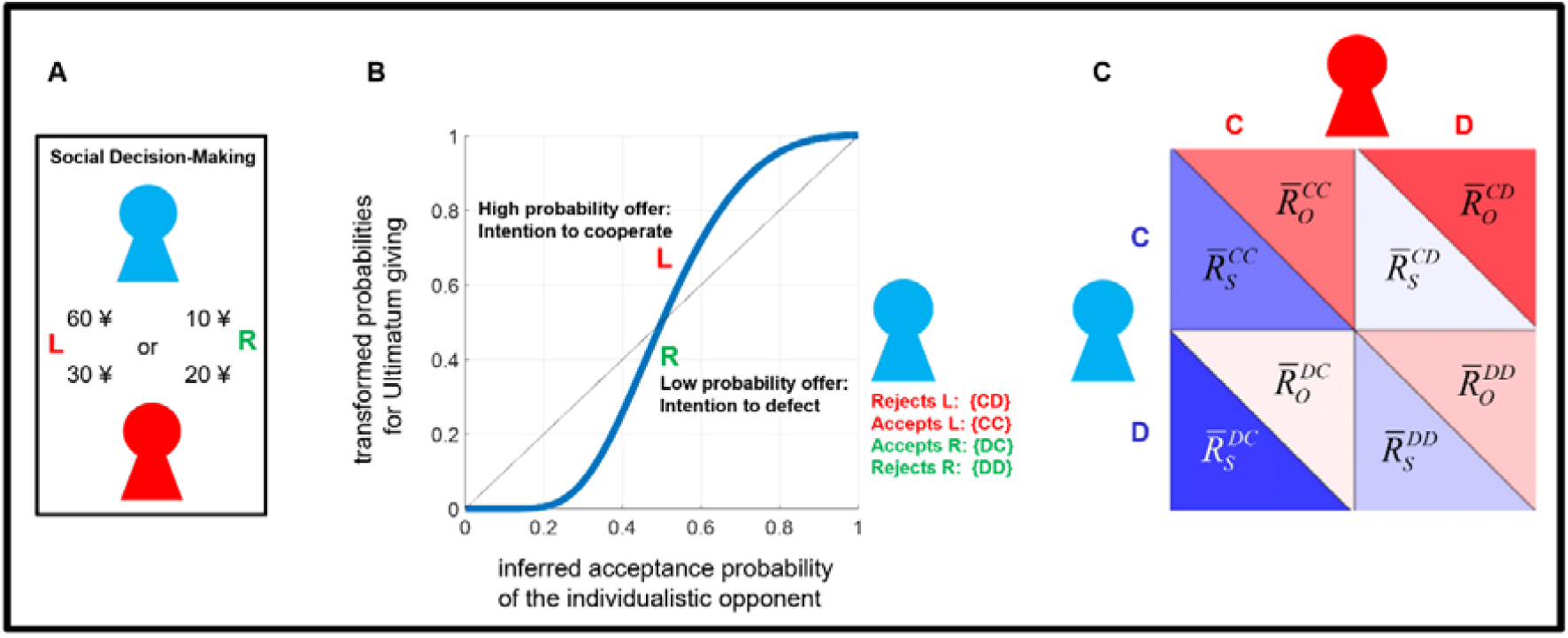
Schematic outline of recovering IPD matrices from an iterative Ultimatum Game. **(A)** Participants (the red icon) chose between two Ultimatums to be given to a computerized opponent (blue icon representing the individualistic opponent) which accepts or rejects them following a probabilistic social value function (see Methods, Eq. 6). At this stage only the participant can see both of the offers (L: left offer; R: right offer). Once selected, the offer is presented to the responder/opponent. The opponent can only evaluate the chosen option but cannot know whether it is better or worse than the unchosen one. The UG included 120 trials in total, that participants played twice (one time against each opponent: individualistic or prosocial). Panel adapted from Pulcu and Haruno (2020). **(B)** Previously, we showed model-based evidence to suggest that participants track their opponents’ acceptance probabilities (on x-axis, also q_A_ in Eq. 6), which are transformed by a nonlinear probability weighting function in the UG. Thick blue line represents the weighted probabilities used while choosing between binary Ultimatums, transforming participant’s inference about the opponent’s acceptance probability. As a result, choosing the offer associated with higher acceptance probability indicates participant’s intention to cooperate with the opponent. Given that opponents make decisions probabilistically following a [social] stochastic value function (q_A_), it is possible to index all outcome types of the PD. **(C)** Schematic diagram outlining the recovered PD matrix from the UG experiment. For example, 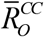 refers to participant’s reward amounts averaged across all conditions satisfying CC outcomes (calculated from the lower rows of reward magnitudes for the chosen option in panel 1A), whereas 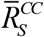 refers to the opponent’s average reward amounts for the same condition. Panel adapted from Press and Dyson (2012).

## Results

### Reformulating an iterative Ultimatum Game as an iterative Prisoner’s Dilemma

If we take Press and Dyson’s original claim at the face value, it should be possible to make quantitative transitions between IPD and UG, or vice versa. Indeed, it is possible to construct PD matrices by indexing trials by: (i) the participants’ inferred choice probability for each of the [computerized] opponents (i.e. *s*∈{*i, p*}, i:individualistic opponent, p: prosocial opponent, also see Methods) generated under the best-fitting computational decision model that accounts for participant choice behaviour^19^, where choosing an option with higher inferred acceptance probability represents the participant’s intention to cooperate: “C”); and (ii) the opponent’s responses to these offers (e.g. stochastically rejecting the offer: “CD”; or accepting the offer: “CC”). After repeating this step for all possible combinations of *xy* ∈{*CC, CD, DC, DD*}, the average reward amounts *R*_*S*_ (i.e. reward to self) and *R*_*O*_ (i.e. reward to opponent) can be calculated across all trials satisfying these conditions, and the cells of the 2x2 matrix for both players can be constructed (Figure 1C). Figure 2A shows an example 2x2 matrix which satisfy the payoff conditions of a PD game.

**Figure 2.**
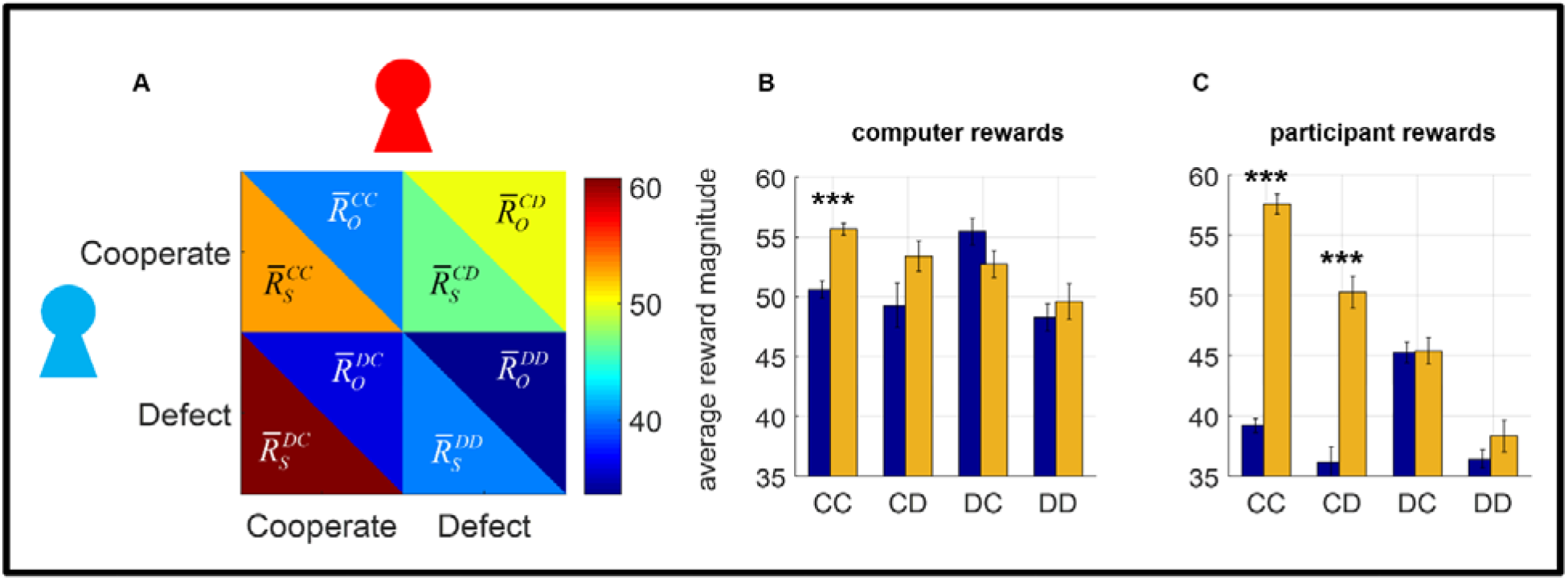
Analyses of payoffs in the IPD. **(A)** An example payoff matrix recovered from the UG experiment satisfying the conditions of the PD game [with an uneven payoff structure]. As in Figure 1C, red icon represents the participant, and the blue icon represents the opponent following an individualistic value function. Color bar represents the average reward magnitude in each of the cells. **(B)** Computerized opponent with prosocial value function (orange bars) is able to acquire significantly higher rewards establishing mutual cooperation with the human participant relative to the individualistic strategy (blue bars, ***p<.001, Bonferroni corrected for multiple comparisons). In line with labels in Figures 1C and 2A, IPD conditions are labeled first with reference to the opponent response (i.e. CD meaning opponent cooperated when human participant defected) **(C)** Human participants acquired significantly higher rewards under mutual cooperation with a prosocial opponent relative to the individualistic opponent both when they cooperated and defected (***p<.001, Bonferroni corrected for multiple comparisons). Error bars designate ±1 SEM.

These transformations demonstrate that it is possible to recover IPD matrices, which are more suitable for evolutionary game-theoretic fitness analysis, directly from an iterative UG experiment. Here, it is important to point out that, although these payoff matrices are introduced with a general reference to IPD, it would be more accurate to define them as “2x2 evolutionary game” matrices which were uniquely constructed for each interaction between human participants and computerized opponents (i.e. 50 nonclinical/healthy volunteers x 2 different computerized opponents (prosocial versus individualistic), in total 100 unique interactions). Press and Dyson (2012) explicitly stated that their formulations apply to all 2x2 evolutionary matrix games, not only those which satisfy the conditions of the Prisoner’s Dilemma Game (e.g. CC>DD).

### Human players cannot escape extortion against individualistic opponents

The next step involves an analysis of the payoff amounts shown in Figure 2B-C. Here, it is important to clarify that cells corresponding to opponent defection (i.e. the cells in the lower row in Figure 2A, based on rejected offers in the UG) actually designate the average amount of forgone rewards for both sides^22^, as both sides get nothing when an offer is rejected in the UG. This is exactly what Press and Dyson referred to in their abstract. In this approach, the average rewards acquired under opponent cooperation (i.e. CC and CD conditions in Figure 2B-C) may be more informative and conservative. The results indicate that opponents with a prosocial value orientation was able to acquire significantly more rewards than the individualistic strategy (based on 2x2 multivariate ANOVA, F(1,98)=15.046, p<.001, Figure 2B). Similarly, the human players were also able to acquire significantly higher rewards playing against a prosocial relative to an individualistic opponent (F(1,98)=297.33, p<.001, Figure 2C). On the other hand, the individualistic opponent acquired significantly more rewards relative to human participants under participants’ unilateral defection (i.e. CD outcomes) or mutual cooperation (comparison of blue bars between Figure 2B and 2C, F(1,98)=98.123, p<.001). However, when it comes to social interactive decision-making with the prosocial opponent, there was a significant outcome by player-type interaction (F(1,98)=4.585, p=.035), whereby human participants were able to acquire higher rewards relative to the prosocial opponent, but only under mutual cooperation (t (98) = 1.979, p=.051). Taken together, these findings indicate that interactions with prosocial opponents lead to mutually rewarding outcomes for human players, whereas the individualistic opponents were able to establish extortionate terms (akin to what ZD strategies can achieve) over the human participants, acquiring significantly higher rewards irrespective of whether the human participants cooperated or defected (i.e. comparison of blue bars for CC and CD conditions between Figure 2B and 2C). It is important to highlight that numerical components of the stimuli (i.e. offer distributions) in the original UG experiment^19^ were identical across all conditions, so the differences reported here cannot be attributed to such factors which might otherwise confound the results (i.e. similar to IPD having a predefined payoff matrix).

### Theory of Mind guided value-based decision-making promotes interpersonal cooperation

In the preceding section, we demonstrated that iterative UG can be reformulated in terms of iterative PD, revealing that human players cannot escape extortion against individualistic opponents. In the next step, it is important to demonstrate how transition probabilities can be computed from the raw data. These describe the rates of cooperation following all possible outcomes *xy* in the IPD. Analyzing 2x2 social interactive decision-making games by this approach is computationally efficient because by constructing Markov transition matrices it is possible to treat these games as a competition between two memory-1 strategies (i.e. describing agent cooperation in terms of the probability with which it changes its behaviour from the previous trial after observing the joint outcome, commonly known as *transition probabilities*). To the best of my knowledge this approach has not been implemented on any data collected from human players.

This approach suggest that the prosocial strategy was significantly more likely to cooperate with the human participants relative to the individualistic strategy (F(1,98)=448.34, p<.001, Figure 3A). This is only reassuring, as the prosocial strategy was designed to follow a social-value function that treats self-other inequality more negatively (see the details in Methods).

**Figure 3.**
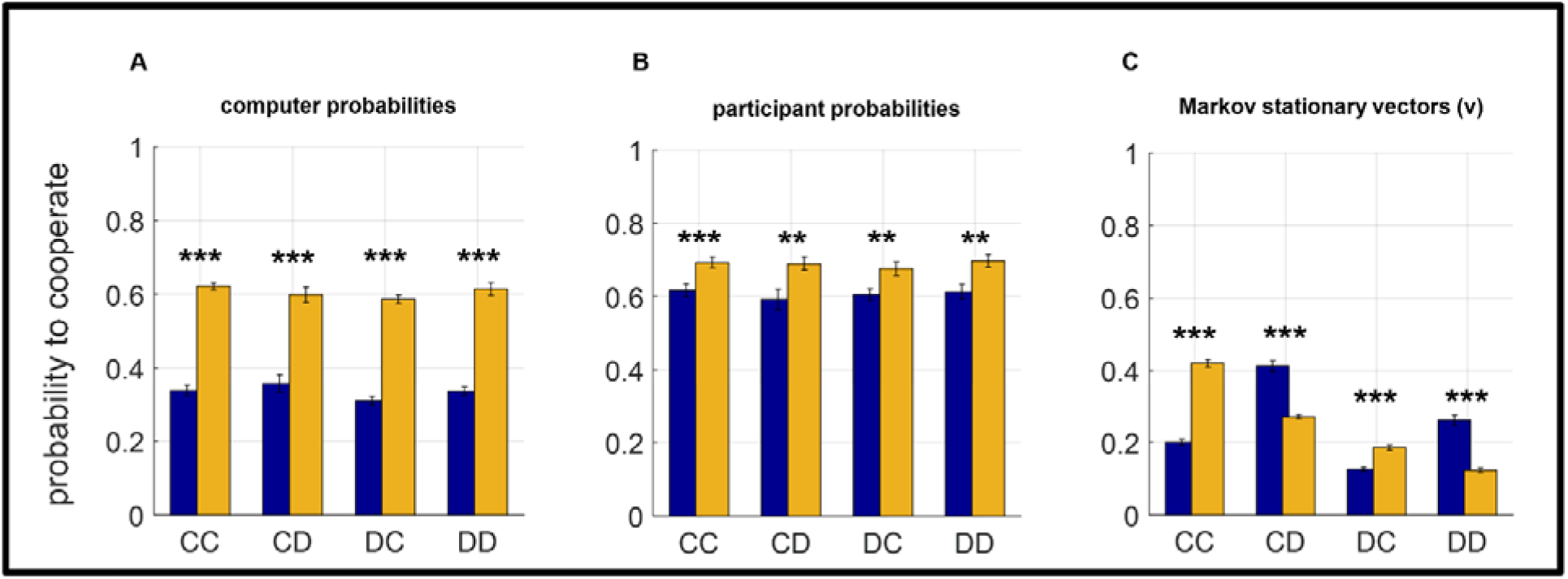
Analyses of transition probabilities in the IPD. **(A)** The prosocial opponent was more likely to cooperate relative to the individualistic opponent after all types of outcomes in the IPD (***p<.001). **(B)** Similarly, participants also cooperated at a significantly higher rate while playing against the prosocial relative to the individualistic opponent (all t (98)>2.91, all p<.005, ***p<.001, **p<.01). **(C)** Summary of Markov stationary vectors computed by Eqs. 2 and 3. All comparisons were statistically significant (***p<.001, Bonferroni corrected for multiple comparisons). Error bars designate ±1 SEM. Orange bars: prosocial opponent; blue bars: individualistic opponent.

In IPD terms, this would translate to a lower probability of defection as unilateral defections lead to self-other inequality in payoffs. Although participants were not explicitly informed about the social value orientation (SVO) of their opponents (i.e. whether the opponent was prosocial or individualistic, they could only infer their opponents’ value functions from a 180 trial observational social-learning session they completed prior to the UG in which they were trying to predict whether opponents would accept or reject offers coming from a third party), they acted in a significantly more cooperative manner while playing against the prosocial opponent relative to the individualistic opponent (F(1,98)=17.295, p<.001, Figure 3B).

Furthermore, although participants were asked to bank as much money as possible in the experiment, which was paid to them in real monetary terms, they acted in a significantly more cooperative manner relative to the computerized opponents (i.e. comparison of blue and orange bars between Figure 3A and 3B, F(1,98)=229.53, p<.001 vs the individualistic opponent; F(1,98)=30.152, p<.001 vs the prosocial opponent). This suggests that ToM driven value-based decisions during social-interactive decision-making (i.e., incorporating opponents’ value functions into one’s own expected value computations) promote interpersonal cooperation. Therefore, interpersonal cooperation emerges naturally as the option with better expected value for human players, irrespective of opponent type.

### Interpersonal cooperation erodes in time

An important follow-up question was asking whether interpersonal cooperation could be sustained. In order to address this question, it was necessary to test the stability of these social interactive systems at the equilibrium point. This involves computing the trajectory of the interpersonal competition from the probabilities of cooperation following any outcome type *xy* in the IPD^23^. Following an identical approach as reported by Press and Dyson and using the vectors of probabilities defining cooperation rate as calculated from the raw data, as shown in Figure 3A and 3B (i.e. two for each participant against each of the opponents (**s** _*p*_, **s**_*i*_); and one for each computerized opponent against the participant (**p***s*, **i***s*)), Markov transition matrices (**M**) can be constructed for each pair accordingly:

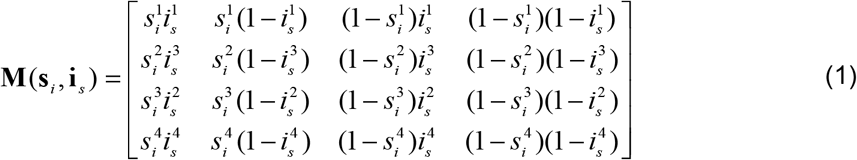

Note that matrix formulated above illustrates an example exchange between a human player and an individualistic opponent. After that the stationary vectors, **v**, of the Markov transition matrices [which satisfy]:

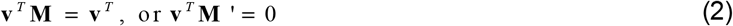

were computed. Markov stationary vectors identify the stable transition probabilities which will be achieved for each pair (e.g. a human participant vs the individualistic opponent), if these interactions continue for a very long time (i.e. the finite evolutionary horizon). These analyses were conducted for each pair (i.e., all combinations of 50 human players vs 2 opponents, 100 pairs in total). Here, it is important to highlight the necessity of this approach as the system stability (as shown in Figure 3C) cannot be predicted from individual players’ transition probabilities (Figure 3A-B). This approach demonstrates several significant results. On one hand, systems involving an interaction with a prosocial opponent was significantly more likely to sustain mutual cooperation (t(98)=16.37, p<.001) and restore cooperation after opponent’s defection (i.e. the DC outcome, t(98)=6.25, p<.001) relative to those which involved an interaction with an individualistic opponent. On the other hand, systems involving interaction with an individualistic opponent were significantly more likely to restore cooperation after participant’s unilateral and both player’s mutual defection (t(98)=8.94, p<.001 and t(98)=8.80, p<.001 for CD and DD outcomes, respectively), relative to those which involved competition with a prosocial opponent.

The results at the stationary state also suggest that interpersonal cooperation will erode in time, considering that cooperation levels at the stationary state (Figure 3C) were globally lower than individual rates calculated from the experiment (Figure 3A-B).

### Behavioural phenotypes that can safeguard long-term cooperation

Erosion of interpersonal cooperation as demonstrated in the preceding section highlights that it is crucial to identify behavioural phenotypes that can safeguard long-term interpersonal cooperation. Using Eqs. 1 and 2 across 2x10^4^ simulations, it is possible to identify [by reverse engineering] the best versus worst strategies in terms of long-term mutual cooperation. After 2x10^4^ simulations, the results described below are robust and fully reproducible even if probabilities stored in the matrix **M** (**s**_i_, **i**_s_) in Eq.1 are generated randomly (e.g. using MATLAB’s *rand* function). An important insight arising from these simulations is that strategies that fare best in terms of maintaining long-term mutual cooperation are in fact the worst when it comes to breaking cycles of mutual defection (Figure 4A), and vice versa. The comparison of individual player profiles (Figure 4B vs C) suggests that players that are projected to maintain high levels of long-term cooperation are globally more cooperative across the board (i.e. following different outcome types CC, CD, DC, DD) and likely to establish mutual cooperation online (i.e. during the interactive decision-making phase). Correlations computed within each group [in order to prevent Simpson’s paradox^24^] suggest that online mutual cooperation (CC) rates are the best predictor of long-term mutual cooperation for best faring players (within top 5% of simulated agent pairs, r=.429, p<.001), whereas online cooperation after DD outcomes correlate significantly with long-term mutual cooperation for the worst faring players (within bottom 5% of simulated agent pairs, r=.151, p<.001). The relationship between each player’s average cooperation rates and mutual cooperation probability at the stationary state suggests a positive trend (Figure 4D; from bottom left to top right increase) albeit with some nonlinearity in the parameter space.

**Figure 4.**
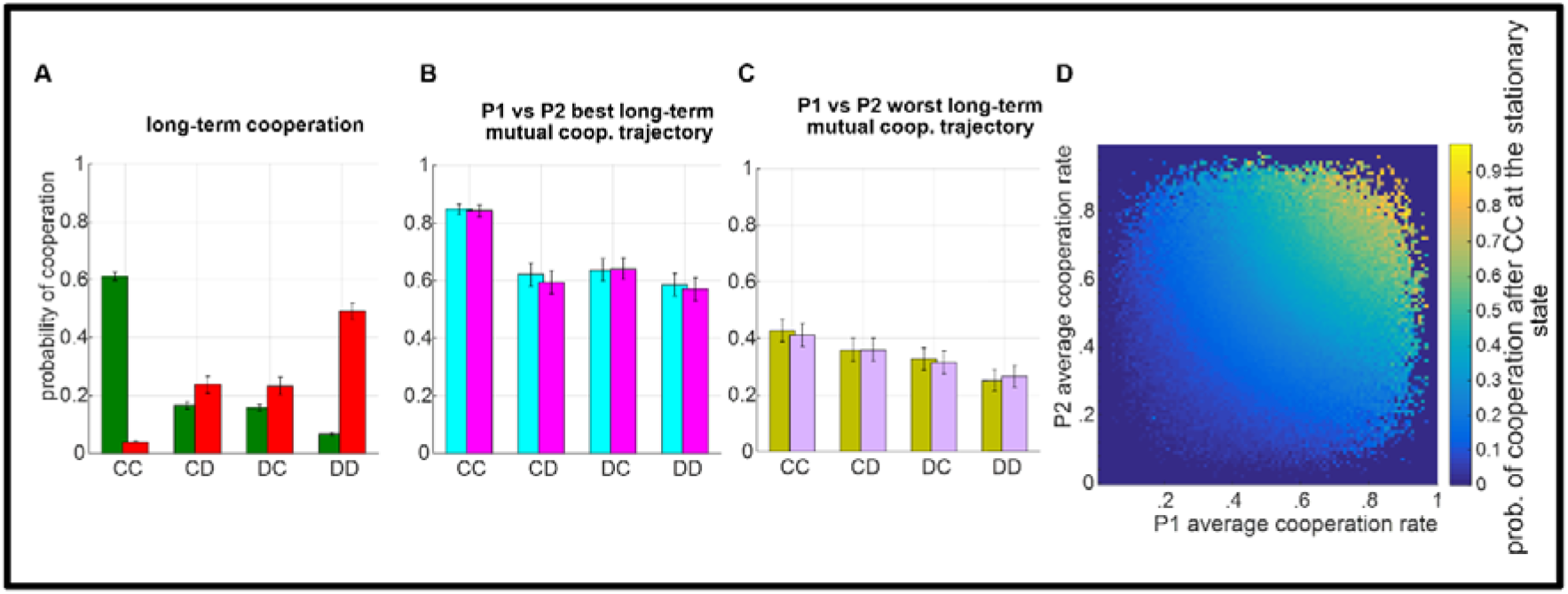
Simulated results for maintaining long-term interpersonal cooperation. **(A)** System stability for best (green bars) and worst (red bars) faring players. Online cooperation rates for **(B)** best and **(C)** worst faring players. 1000 simulated agents per group. Error bars designate ±1 SEM. **(D)** The relationship between P1 vs P2 average cooperation rates across all outcome types and long-term mutual cooperation at the stationary state. Dark blue regions in the parameter space is not covered due to average cooperation rates not pushing to the extreme values.

### Human behavioural phenotype risks extinction against individualistic strategies

After computing the transition probabilities at Markov stationary states by Eq.2, it is relative straightforward to calculate each player’s expected outcome (i.e. evolutionary fitness). The expected payoff of any agent at the stationary state is computed by

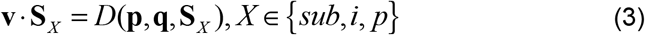

Where 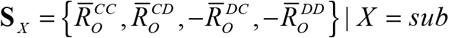, etc. contains the numerical reward values of each agent’s unique 2x2 matrix in a vector format (Figure 2B and 2C); while **p** and **q** replace each agent’s vector of cooperation probabilities which were previously described (i.e. **s**_*i*_ and **i**_*s*_). Here, *D*(**p**,**q**,**S**_*X*_) requires no normalisation considering (**1**· **v**) = 1. Note that average reward amounts for DC and DD outcomes are negated in this calculation as these represent forgone reward amounts due to opponent’s rejection in the UG (i.e. the reward amounts sacrificed to convey that the proposed offer amount was *unfair*). Each player’s evolutionary fitness at the Markov stationary state were computed, revealing how human players would fare against their opponents (Figure 5). The analysis of evolutionary fitness by a 2x2 (opponent type (individualistic vs prosocial) x agent type (computer vs human)) multivariate ANOVA suggested a main effect of opponent type (F(1,98) =80.377, p<.001) and a marginally significant opponent x agent type interaction (F(1,98)= 3.805, p=.054). Here, the significant main effect of opponent type highlights the accumulated resource gap between individualistic and prosocial agents, considering that numerical values of the task environment was identical across all pairwise interactions. Although subsequent pairwise comparisons between human participants and computerized opponents were not statistically significant (all p>.105), the results from the Markov stationary states indicate that humans would prevail over prosocial strategies (extinction rate: *p*(**v** · **S** _*p*_ > **v** · **S** _*sub*_) = 0.14), whereas they would fare mostly poorly against individualistic strategies (*p*(**v** · **S***i* > **v** ·**S***sub*) = 0.70, also see Figure 4). This would be in line with the preceding results suggesting that human players fail to escape extortion against individualistic opponents.

**Figure 5.**
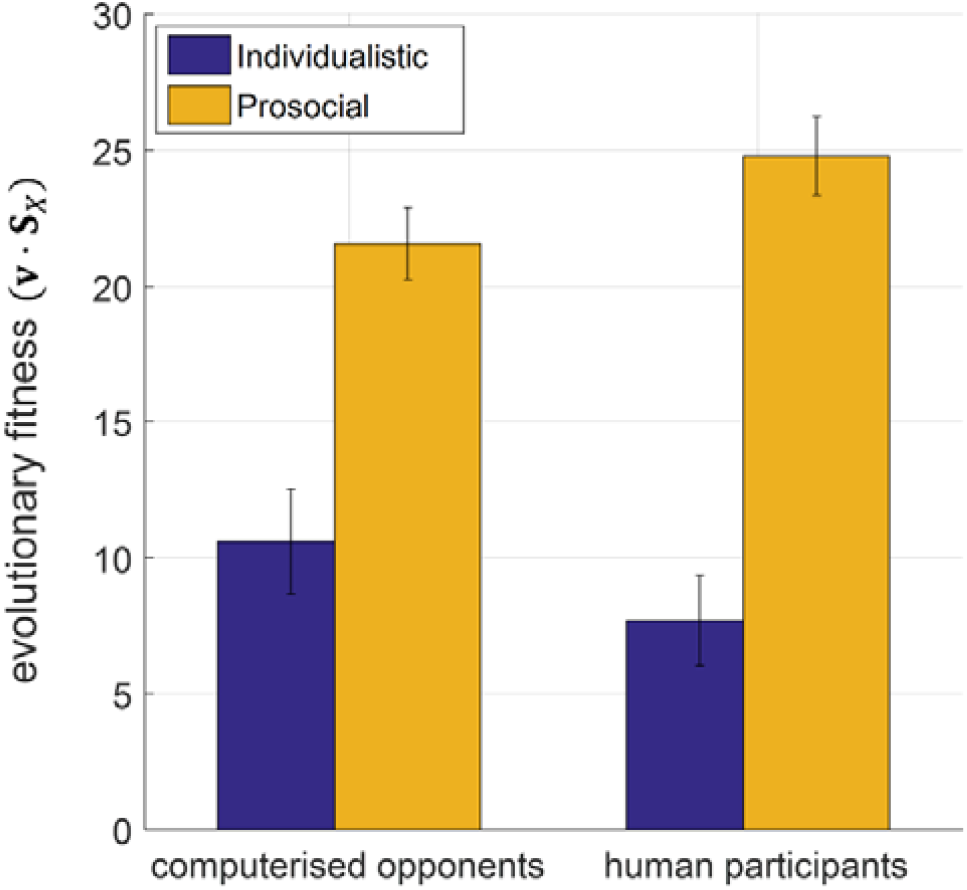
Evolutionary fitness at the Markov stationary states. Irrespective of agent type (i.e. human players or [computerized] opponents) evolutionary fitness is significantly higher in social interactions with a prosocial opponent (main effect of opponent type, ***p<.001). Error bars designate ±1 SEM.

For completeness, note that human players’ total evolutionary fitness (*f*) can be calculated as:

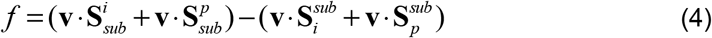

### Do zero determinant strategies exist in the wild?

In the preceding sections, the analysis of average payoff amounts (Figure 2B-C) suggested that individualistic opponents can establish extortionate terms over the human players, which is one of the key properties of the ZD strategies discovered by Press and Dyson (2012). Although Press and Dyson’s (2012) discovery posits the existence of ZD strategies algebraically, this does not necessarily mean that they emerge naturally during social interactive decision-making between two competitors. To estimate the probability with which ZD strategies can be observed in a typical lab-based social interactive decision-making experiment, we investigated whether a direct linear relationship is established between the payoffs of the two players (i.e. opponents vs human players) in our recruitment cohort. Note that previous studies implementing the ZD framework to human decision-making in the IPD^25^ approximated these linear relationships between payoffs by relying on correlations rather than trying to enforce necessary algebraic conditions in which the determinants would vanish.

Following Press and Dyson’s notation (also see their Eq. (7)), I investigated such possibility, 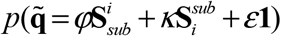, by numerically estimating the values of *φ,κ,ε* (using MATLAB’s *fmincon*) which maximized the function:

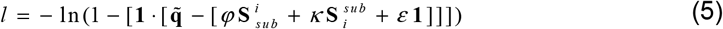

and computed *p*(*l*) = 0 across all interactions in our recruitment cohort (computerised opponent vs human player and vice versa). If *p*(*l*) = 0 is satisfied, it would mean that the determinant vanishes and it would suggest that [in this example, opponent’s] strategy is truly a ZD strategy against the human player. However, out of 100 combinations between computerized opponents and human players it was not possible to identify any strategy truly satisfying this condition.

## Discussion

The majority of human experimental studies investigating interpersonal cooperation by using variants of the PD game^26-28^ used well-established, yet somewhat rigid, computerized strategies such as *tit-for-tat* or *tit-for-two-tats*, which were previously thought to be evolutionarily competitive^5^. The main reasoning behind using these strategies in behavioural experiments is to understand how human participants fare against strategies which are shown to have competitive evolutionary fitness, as established by theoretic simulation-based studies. Later expansions of the IPD strategy landscape by Pavlovian^29^ and ZD strategies^14^ provided highly novel but mostly theoretical insights about evolution of prosocial behaviors^2^. The current experimental approach, founded in bridging two key behavioural economic paradigms, provides novel insights about human interpersonal cooperation particularly against agents that follow a stochastic social-value function, just as human decision-makers do in behavioural economic experiments^19^.

By following up Press and Dyson’s (2012) proposal that a convergence between IPD and UG can naturally emerge when a player with ToM engages in social interactive decision-making, I provided quantitative evidence in favour of this proposal, which essentially transforms human Ultimatum bargaining behaviour to be more suitable for analysis using an evolutionary game theoretic approach. The results communicated in this work suggest that ToM sophistication often exhibited by human players^15^ naturally promotes interpersonal cooperation even when players are self-interested, trying to maximise their short-term monetary gains (Figure 3). However, the current results also suggest that social value orientation (SVO) of the opponents play a significant role, and human players cannot escape extortion by opponents with an individualistic social value orientation (Figure 2B-C), to such degree that they would eventually risk extinction more frequently than not. Irrespective of opponents’ SVO, equilibria at the stationary states for these social interactive systems suggest that interpersonal cooperation will erode substantially in time (Figure 3C).

Exploratory simulations complementing these findings suggested that maintaining online mutual cooperation is the strongest predictor of long-term mutual cooperation, whereas ability to break cycles of mutual defection (i.e. higher probability of cooperating after DD outcomes) was associated with poor long-term mutual cooperation rates. This is also similar to how individualistic agents behaved in the experiment, overall performing poorly in terms of their reward trajectory (Figure 5), yet driving cooperative human players (Figure 3B) to [near] extinction.

Results communicated in this work suggest that opponents’ SVO (i.e. individualistic versus prosocial) directly influence players’ payoffs. For example, sustaining mutual cooperation with prosocial agents yield much better payoffs for human players. At the same time, human players were shown to give in to extortion by individualistic opponents (Figure 2B-C). Despite following very different methodologies, these findings are strikingly in line with the results of a previous IPD study in which healthy volunteers played against different computerized agents following stationary conditional probabilistic strategies^22^. Subsequent work suggested that extortionate strategies (i.e., similar to the opponent with individualistic SVO) can outperform generous (i.e. similar to the opponent with prosocial SVO) strategies that play against human players, but only if the experiment duration is reasonably long^25^ (500 trials). In our experiment (across 120 trials), human participants tried to bank as much money as possible^19^ and they ended up acting in a significantly more cooperative manner than their opponents while doing so (Figure 3A-B). A subsequent evolutionary stability analysis at the Markov stationary states (Fig.3C) revealed that systems involving interactions with prosocial opponents are significantly more likely to sustain mutual cooperation (i.e. CC outcomes), whereas systems involving interactions with individualistic opponents are more likely to break cycles of mutual defection (i.e. DD outcomes), albeit this maybe at the expense of human participants settling down for less (Fig. 2C). Over finite evolutionary horizons these differences converge and show that human participants would prevail over prosocial, but fare poorly against individualistic agents that acted in an extortionate manner (Fig. 5). This is also in line with findings from previous studies that used longer behavioural protocols^25^. Taken together, these findings suggest that human players focusing on maximizing short-term goals may be oblivious to, and unprepared for the long-term consequences of their decision strategies. This shortsightedness would undermine long-term evolutionary/social-economic fitness in a world in which humans need to interact with artificial agents or even corporate entities which should be quantitatively better equipped to estimate their reward trajectories^30^.

Only a limited number of studies investigated interactions between human players and ZD strategies. These studies formulated ZD strategies in terms stationary conditional probabilities derived such that there would be a correlation between the computerized agents’ and the human players’ payoffs^22,25^, allowing extortionate ZD strategies to acquire higher, whereas generous ZD strategies to acquire lower rewards. This approach satisfies the linear relationship prerequisite defining ZD strategies originally formulated by Press and Dyson. In the current work, I took a different approach and derived the conditional probabilities of the computerized agents from their stochastic decisions to accept or reject Ultimatums. As a result, the conditional probabilities of the computerized opponents were varied in the current work for each participant (as these also depended on participant choice behaviour, Fig. 3A-B) and each opponent, allowing this work to account for wider individual variability and establish a birds-eye view on the long-term trajectory of human interpersonal cooperation. Within this inherent variability in participant and opponent strategies, I numerically estimated the degree to which a stringent linear relationship can be established between the payoffs of humans versus computerized opponents (both players following a stochastic social value function), as mathematically formulated by Press and Dyson 2012, and vice versa. Under these conditions, the estimated likelihood of ZD strategies emerging naturally in cooperative/competitive interactions would be <1%. However, considering the extent of biodiversity on planet Earth and the vast number of interpersonal exchanges taking place between billions of people every day, I do not think these results undermine the ecological validity of zero determinant strategies or the influence of this framework.

To the best of my knowledge, the current work is the first to follow up Press and Dyson’s hypothesis of expressing UG in terms of IPD. Here, I demonstrate that in interactions between human participants with ToM sophistication playing against opponents following a stochastic social value function, it is possible to bridge these social economic games. Although the wide majority of the previous literature focused on human responder behaviour in the UG^9,12,28,31,32^, the present manuscript based on our recent work describing value computations underlying human proposer behaviour^19^, demonstrates that more studies investigating this aspect of social interactive decision-making can reveal rich and multidimensional insights about the nature and mechanisms of human prosocial behaviors.

## Materials and Methods

### Participants

Fifty nonclinical volunteers were recruited for the Ultimatum giving experiment. Initially, participants were asked to learn the preferences of two computerised opponents in an observational learning session. Opponent strategies were based on social-value functions (q_A_) that generated online accept or reject decisions from self-versus-other reward distributions probabilistically, globally agreeing with a prosocial and an individualistic social value orientation (SVO^20^):

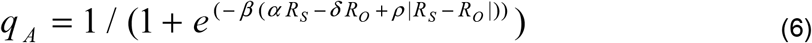

In the formula above, R_S_ refers to the share of the reward for the computerized agent, whereas R_O_ refers to the reward amount to the participant (Fig. 1C). The hyper-parameters defining the valuation of the agents [α, δ, ρ, β] were set to [1.096, 0.382,-2.512, 0.037] for the individualistic agent; and [1.368,-0.644,-3.798, 0.045] for the prosocial agent.

In the subsequent step after adequate learning, participants were asked to choose between two monetary Ultimatums to be given to these opponents that makes “accept” or “reject” decisions following the stochastic social-value function shown in Eq.6 (also see Fig.1A, experiment involving 120 binary decisions against each opponent type). The results from the original experiments were published previously^17^ and the current analytic approach concerns the reanalysis of that dataset readily available at : https://osf.io/qy9e6/?view_onlycbd3eac3cd0744f69ba0828033789e61

